# Multi-scale microenvironment covariance for unsupervised spatial domain identification

**DOI:** 10.64898/2026.10.20.760373

**Authors:** Bairui Du, Hisanori Kiryu

## Abstract

**Motivation:** Many spatial domain identification methods describe the neighbourhood of a cell by mean gene expression. However, the mean cannot separate neighbourhoods that differ only in gene–gene covariance and tissue is organised at several spatial scales.ht

**Results:** We propose CoSMIM, which represents the niche of each cell by a weighted fusion of gene–gene covariance matrices of its spatial neighbourhood at multiple scales. Principal component analysis and Leiden clustering of the matrix square root of the fused matrix identify spatial domains without model training. On six benchmark datasets from five platforms, CoSMIM attains the best overall rank over eight evaluation metrics among 16 methods. It also ranks first on a Xenium mouse brain dataset, and on all seven datasets its median NMI exceeds that of single-scale COVET. The domain covariance matrices also show gene pairs whose correlation differs between domains.

**Availability and implementation:** The source code, documentation and test data of CoSMIM are available at GitHub (https://github.com/barondu/cosmim).

**Supplementary information:** Supplementary data are available at *Bioinformatics* online.

## 1. Introduction

Tissues are organised into spatial domains, for example the layers of the cerebral cortex and organ primordia such as the liver bud in the early mouse embryo. Spatial transcriptomics measures gene expression with spatial coordinates, allowing these domains to be reconstructed at the molecular level (Rao et al., 2021; Moses and Pachter, 2022). Partitioning a tissue into transcriptomically coherent, spatially contiguous regions is therefore the common entry point to tissue annotation, microenvironment–phenotype association and developmental trajectory mapping.

Cells belong to the same spatial domain because they share a microenvironment: the neighbouring cells, signalling molecules and local tissue architecture around them. One domain may contain several cell types, so domain identification clusters microenvironments rather than cell type identities (Fig. 1b). We model it as a local neighbourhood around each focal cell, its niche. In turn, the niche can affect the transcriptional state of the cell. Previous studies have attributed a substantial part of the expression variation within a cell type to the composition of the niche (Fischer et al., 2023; Agrawal et al., 2024).

**Fig. 1:**
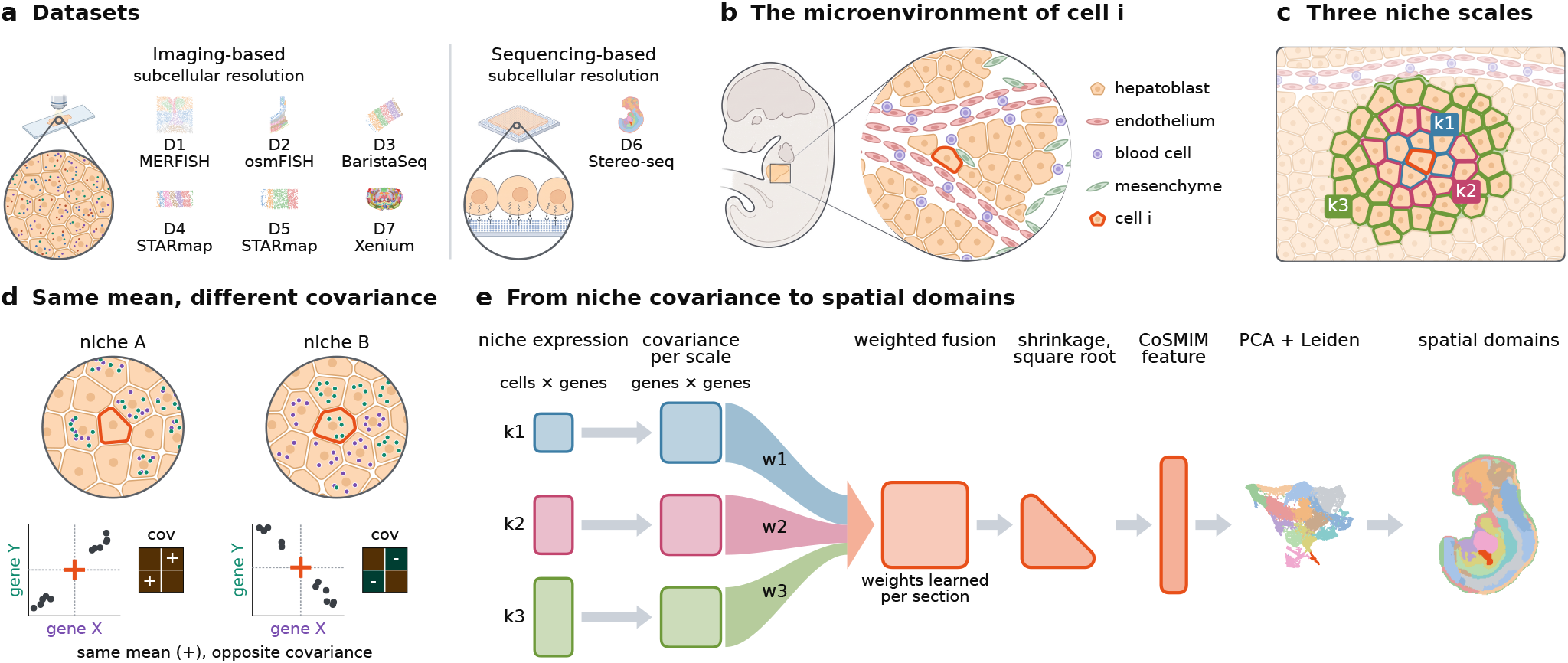
Overview of CoSMIM. **(a)** The seven datasets D1 to D7 and their platforms. **(b)** The microenvironment of a cell *i* in an embryonic liver contains several cell types. **(c)** The niches of cell *i* at three scales: cell *i* and its *k*_1_, *k*_2_ and *k*_3_ nearest neighbours. **(d)** Genes X and Y have the same mean expression in niches A and B, but their covariance is positive in A and negative in B. **(e)** At each scale, the niche expression (cells *×* genes) gives a gene–gene covariance matrix. The three matrices are fused with weights *w*_1_, *w*_2_ and *w*_3_, which are learned for each section. The fused matrix is shrunk, and its matrix square root gives the CoSMIM feature. PCA and Leiden clustering of this feature give the spatial domains. The drawings in (a) and (b) were created in BioRender (Du, 2026).

We call the mean expression of each gene over a niche a first-order statistic and the gene–gene covariance over the niche a second-order statistic. Here, order refers to the statistical moment, not to neighbours in a spatial graph. Two niches can have the same mean expression profile but different gene–gene covariance (Fig. 1d). This covariance can reflect cell-type co-occurrence, cell– cell interaction and co-regulatory programmes (Haviv et al., 2025; Acharyya et al., 2022; Du and Kiryu, 2025). A first-order statistic has one value per gene and none per gene pair, so it cannot describe how the correlation of two genes differs between niches.

In addition, tissue is organised at several spatial scales, and the number of neighbours *k* sets the scale that a niche describes (Fig. 1c). These scales range from direct contact between cells (Prevot et al., 2010), through the cell types that share one cortical layer (Tasic et al., 2018; Codeluppi et al., 2018), to a tissue region such as the mesenchyme around the liver bud (Gordillo et al., 2015). A niche defined with a single *k* describes mainly one of these scales.

Among training-free methods, NichePCA (Schaub et al., 2025) and BANKSY (Singhal et al., 2024) use first-order statistics, and COVET (Haviv et al., 2025) uses a second-order statistic at a single scale (§2.2). To our knowledge, no neighbourhood-based method combines second-order statistics with several spatial scales.

We propose CoSMIM (**Co**variance of **S**patial **M**ulti-scale **I**ntegrated **M**icroenvironment), which combines both properties. CoSMIM computes the gene–gene covariance matrix of the neighbourhood of each cell at multiple scales, fuses them into one matrix with weights, and uses the matrix square root of the fused matrix as a Euclidean feature. In COVET the expression in each niche is centred on the mean of the whole section. As a result, each matrix also contains a first-order term. Standard PCA and Leiden clustering are then applied to this feature, so that no model training is required (Fig. 1). In addition, because the feature is a covariance matrix, the gene–gene correlations within each identified domain can be examined directly.

We evaluate CoSMIM for domain-identification accuracy (§3.1) and for biological interpretation. Its per-domain covariance matrices show gene pairs whose correlation has opposite signs in two domains, the third ventricle and the fornix (§3.2). On one section of the mouse embryo, CoSMIM is the only one of five methods that resolves the liver (§3.3).

## 2. Materials and Methods

### 2.1. CoSMIM

CoSMIM defines a cell’s niche not as its neighbourhood mean expression (first order) but as the set of neighbourhood gene– gene covariances 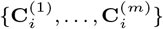 taken over *m* spatial scales *K* = {*k*_1_ *<* · · · *< k*_*m*_} (second order), where 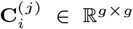 records which gene pairs act in concert (positive) or in opposition (negative) among cell *i* and its *k*_*j*_ nearest neighbours. These matrices are fused with weights into a single *g* × *g* matrix. Here *m* = 3 and K = {5, 15, 45} on all seven datasets. The feature of cell *i* is

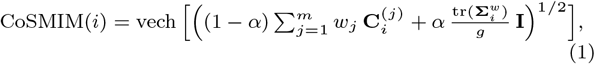

where 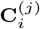 is the neighbourhood covariance at scale *k*_*j*_ (Eq. 2), *w*_*j*_ the scale weights (§2.1.1), 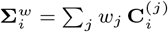the fused covariance, *α* the shrinkage strength (§2.1.2), and vech the lower-triangular half of the symmetric result.

Expression is library-size normalised and log(1+*x*) transformed, and the *g* genes carried into the covariance are those that pass a spatially variable filter and are then taken as highly variable, capped at 64 by default; the quality-control thresholds and the per-dataset values are in Supplementary Data S1. Let the log-expression of each cell after preprocessing be **x**_*i*_ *∈* ℝ^*g*^ . A BallTree on the spatial coordinates retrieves the *k*_*j*_ nearest neighbours of cell *i*, the expression profiles of cell *i* and these neighbours giving the neighbourhood expression 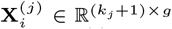. We use COVET-style global mean centring 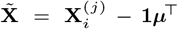(where 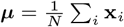_*i*_, making different niches comparable), so that

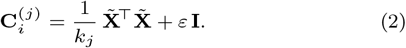

The ridge *ε***I** keeps the matrix invertible when the niche has fewer cells than genes (Supplementary Note S1).

#### 2.1.1. Scale fusion and scale weights

The scale weights *w*_*j*_ of Eq. 1 are learned once for each section by minimising

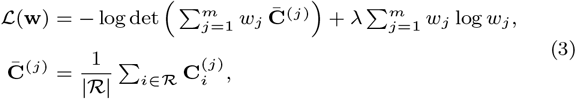

over the simplex *w*_*j*_ ≥ 0, Σ*w*_*j*_*=*1 _*j*_ *w*_*j*_ = 1, where *ℛ* is a fixed random subset of at most 1000 cells of the section. The entropy, term moves the weights towards uniform. Its strength *λ* is the only parameter of the fusion step, and it is set for each dataset (Supplementary Data S1). Supplementary Note S1 gives the solver and the gradient.

#### 2.1.2. Shrinkage and matrix square root

The largest niche neighbourhood scale *k* has 46 cells. On five of the seven datasets the number of genes is larger than this, so the fused covariance is rank deficient before the ridge is added. We therefore shrink the fused matrix towards a scaled identity matrix (Ledoit and Wolf, 2004), with a fixed strength *α* = 0.1 on all datasets. The result is the CoSMIM matrix **Σ**_*i*_. We take its symmetric matrix square root 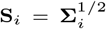 and keep the entries on and below the diagonal as the feature vector (Fig. 1e). The Frobenius distance *∥***S**_*i*_ − **S**_*l*_*∥*_*F*_ is an upper bound on the Bures–Wasserstein distance between **Σ**_*i*_ and **Σ**_*l*_ (Bhatia et al., 2019), and the two are equal when the matrices commute (Supplementary Note S1). As in COVET (Haviv et al., 2025), we therefore use Euclidean distances between the square-root features instead of the Bures–Wasserstein distance, so standard PCA and Leiden clustering can be applied without change. Supplementary Note S3 tests the number and the sizes of the scales and compares the square root with other maps (Supplementary Tables S2–S4 and Supplementary Fig. S1).

### 2.2. Previous niche spatial clustering methods

NichePCA, BANKSY and COVET cluster a feature that is built from the spatial neighbourhood of each cell, as CoSMIM does. We re-run these three methods and a non-spatial baseline, PCA followed by Leiden clustering (PCA+Leiden), with the protocol of §2.5. NichePCA (Schaub et al., 2025) represents each cell by the mean expression of its neighbourhood. BANKSY (Singhal et al., 2024) combines each cell’s own expression with the mean expression and azimuthal Gabor-filtered features of its neighbourhood. COVET (Haviv et al., 2025) characterises the niche by its neighbourhood gene–gene covariance at a single fixed neighbourhood scale. COVET and CoSMIM also differ in shrinkage, gene selection and clustering settings, so the comparison between them reflects these differences together with the number and sizes of the scales (§3.1). MENDER (Yuan, 2024) uses neighbourhoods at several ranges, but it describes them by cell-type composition and not by gene–gene covariance. Deep-learning methods such as SpaGCN (Hu et al., 2021), STAGATE (Dong and Zhang, 2022) and GraphST (Long et al., 2023b) require model training and tuning, and their embeddings are difficult to interpret biologically. SCAN-IT (Cang et al., 2021) treats domain identification as image segmentation and solves it with a graph neural network.

### 2.3. Datasets

We use the public spatial-domain-identification benchmark adopted by NichePCA (Schaub et al., 2025) and SDMBench (Yuan et al., 2024), covering 6 platforms, 7 datasets and 25 sections (Supplementary Note S2). Five of the datasets are small sections from imaging platforms at single-cell resolution, with restricted gene panels and domains that are adjacent nuclei or graded cortical layers: mouse hypothalamic preoptic region by MERFISH (dataset1) (Moffitt et al., 2018), mouse somatosensory cortex by osmFISH (dataset2) (Codeluppi et al., 2018), mouse visual cortex by BaristaSeq *in situ* sequencing (dataset3) (Long et al., 2023a), and mouse visual cortex and medial prefrontal cortex by STARmap (dataset4, dataset5) (Wang et al., 2018). The Stereo-seq whole-transcriptome mouse embryo (dataset6) (Chen et al., 2022) spans E9.5 and E10.5, and many of its organs are alike in mean expression while differing in co-regulatory structure (§3.3). The Xenium mouse brain (dataset7) (Bhuva et al., 2024) has the most cells per section and the most annotated domains and we use it to evaluate CoSMIM on large, finely divided sections. For dataset1 to dataset6 we take the annotations from the same sources as SDMBench. For dataset7, we use the region annotation of Bhuva et al. (2024), who registered each section to the Allen mouse brain atlas. Supplementary Table S1 also compares the methods on dataset4 against the smoothed manual annotation of Singhal et al. (2024).

### 2.4. Evaluation metrics

We use nine metrics in three groups (Supplementary Note S4). The accuracy group compares the predicted domains with the annotation: normalised mutual information (NMI), homogeneity (HOM), completeness (COM) and the adjusted Rand index (ARI). The continuity group measures, without the annotation, whether each predicted domain is spatially contiguous: spatial chaos (CHAOS), the percentage of abnormal spots (PAS) and the average silhouette width (ASW) on the spatial coordinates. The marker group, Moran’s *I* and Geary’s *C*, measures the spatial autocorrelation of the marker genes of each predicted domain. As in SDMBench (Yuan et al., 2024), the methods are ranked within each metric so that the largest rank is the best. SDMBench does not report ARI, so the overall score is the sum of the ranks on the other eight metrics, each computed on the across-dataset medians (Fig. 2d).

**Fig. 2:**
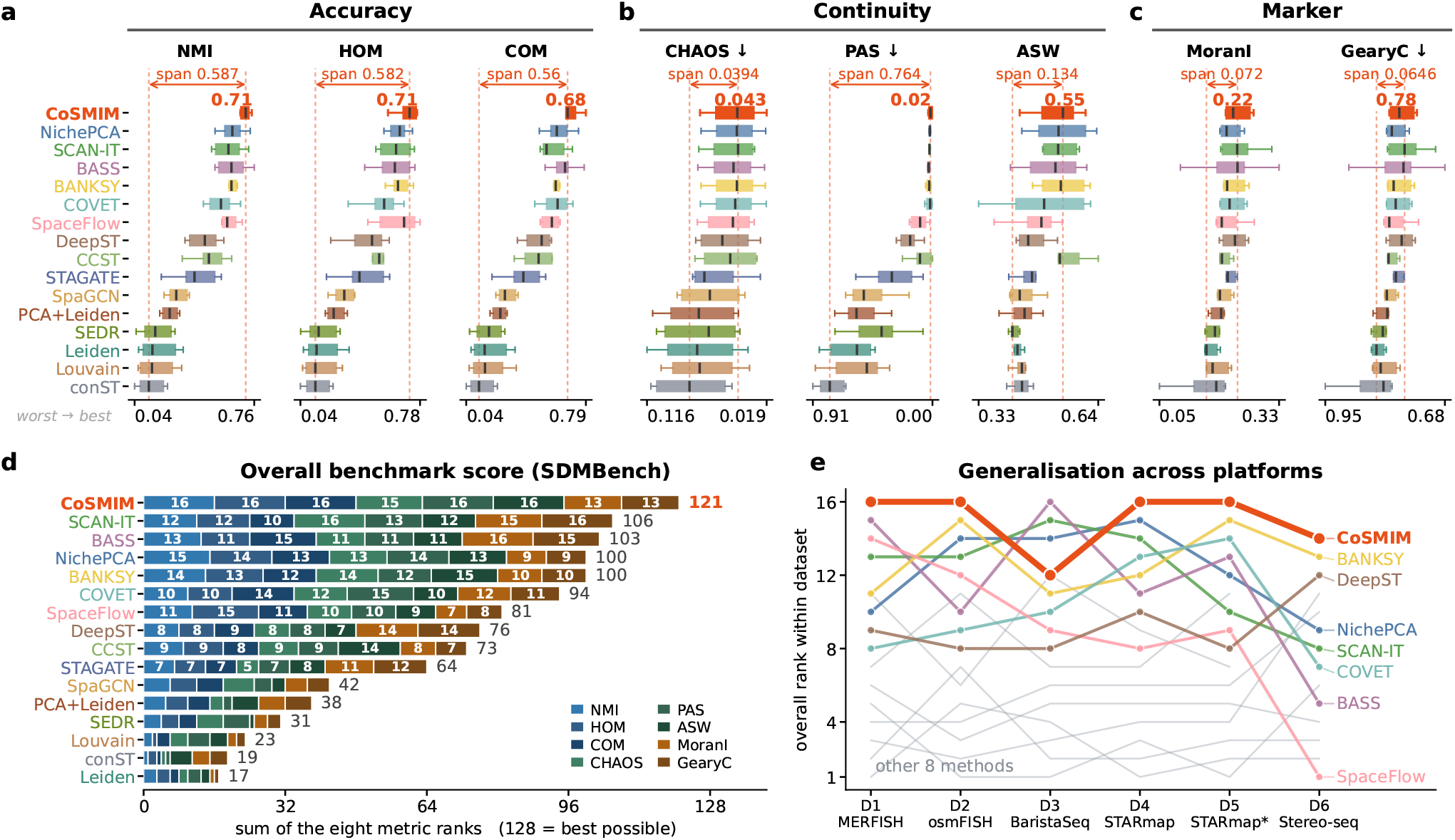
Benchmarking of 16 spatial domain identification methods on six datasets (22 sections). **(a–c)** The eight metrics in three groups: accuracy (a), continuity (b) and marker (c). Each metric is shown on its own value scale as Tukey boxplots, and the line in each box marks the median. The numbers at the two ends of an axis are the extremes of the plotted range. The dashed rules and the arrow above them mark the span of the sixteen methods’ medians. Metrics marked *↓* are better when lower and have reversed axes, so better is to the right throughout. The red numeral is CoSMIM’s median. **(d)** Overall benchmark score (summed rank). This is the sum of the eight metric ranks, each computed on the across-dataset medians (maximum 128). **(e)** Overall rank within each dataset. A larger rank is better. Grey lines show the eight methods with the lowest scores in (d). D1–D6 are datasets 1–6 (§2.3)

### 2.5. Benchmarking protocol and statistical tests

For every method that we run, the features are reduced by PCA and clustered with Leiden (Traag et al., 2019) in Scanpy (Wolf et al., 2018). The Leiden resolution is searched for the number of clusters closest to the number of annotated domains in each section, or to 60 on dataset7 as in Schaub et al. (2025). On dataset7 we cluster each whole section but evaluate only its left hemisphere, because the coronal sections are approximately left– right symmetric. UMAP (McInnes et al., 2018) is used only for visualisation. The primary metric is the median NMI over the sections of each dataset.

On dataset1 to dataset6, SDMBench (Yuan et al., 2024) reports results for 11 of its 13 methods: BASS (Li and Zhou, 2022), CCST (Li et al., 2022), conST (Zong et al., 2022), DeepST (later published as GraphST) (Long et al., 2023b), Leiden, Louvain (Blondel et al., 2008), SCAN-IT, SEDR (Xu et al., 2024), SpaceFlow (Ren et al., 2022), SpaGCN and STAGATE. We place these published results alongside our results for CoSMIM, NichePCA, BANKSY, COVET and PCA+Leiden, after rescaling the published ASW values in the same way as ours (Supplementary Note S4). NichePCA, BANKSY and COVET each have a neighbour count *k*. We ran each at *k ∈* {5, 8, 10, 15, 20}. For each method we selected one global *k*, the value with the best mean rank over NMI, HOM and COM on dataset1 to dataset6 (Supplementary Fig. S7). Each competitor uses this *k* on all seven datasets. CoSMIM uses the same scale set on all seven datasets, but its gene selection, gene cap, number of principal components, number of clustering-graph neighbours and *λ* are set per dataset (Supplementary Data S1).

To compare CoSMIM with the four re-run competitors on the 25 sections, we use two nonparametric tests (Demšar, 2006).

A two-sided paired Wilcoxon signed-rank test (Wilcoxon, 1945) compares CoSMIM with each competitor, with a Bonferroni correction for the four comparisons. The Friedman test (Friedman, 1937) compares the five methods jointly, and the Nemenyi post-hoc test then identifies which pairs differ (Supplementary Note S5).

## 3. Results

### 3.1. CoSMIM gives the best domain-identification accuracy among 16 methods

We first evaluate whether the multi-scale covariance feature identifies spatial domains more accurately than existing methods. Fig. 2 ranks CoSMIM against 15 other methods on dataset1 to dataset6 (22 sections) under the protocol of §2.5. CoSMIM’s summed rank over the eight metrics is 121 out of 128, 15 higher than the second method, SCAN-IT, and 21 higher than NichePCA, the best of the four competitors we reproduced ourselves (Fig. 2d). CoSMIM is first on the three accuracy metrics of Fig. 2 (NMI, HOM and COM), on PAS and on ASW. In addition, CoSMIM ranks first on the 5 datasets other than dataset3 (Fig. 2e). CoSMIM’s lowest-placed group is the marker score, where it places fourth. A possible reason is that CoSMIM defines domains by the neighbourhood gene covariance, whereas Moran’s *I* and Geary’s *C* measure the spatial autocorrelation of individual marker genes. Therefore, to assess the biological significance of the identified domains, §3.2 and §3.3 examine the gene–gene correlations within the CoSMIM domains.

Supplementary Fig. S2 shows the metrics and the spatial domains for each of the seven datasets separately. Of the five methods that we ran, CoSMIM has the highest median NMI on every dataset (Supplementary Data S1). On dataset7, the largest dataset, CoSMIM has the highest NMI on the left hemisphere of each of the three sections. Its median NMI is 0.659, against 0.641 for COVET, 0.637 for NichePCA, 0.614 for BANKSY and 0.455 for PCA+Leiden (Supplementary Figs. S5 and S6).

In addition, we test the NMI differences between CoSMIM and the four reproduced competitors on all 25 sections of the seven datasets. Under the two-sided paired Wilcoxon signed-rank test, CoSMIM has a significantly higher NMI than each of the four competitors (*P* ≤ 9.1 × 10^−4^). After Bonferroni correction for the four comparisons, all four *P* values are still below 0.05. The Friedman test followed by the Nemenyi post-hoc test also separates CoSMIM from each of the four competitors (*P* ≤ 0.044, Supplementary Note S5). NichePCA and BANKSY build their features from first-order statistics. The azimuthal Gabor-filtered features that BANKSY adds also give one value per gene. CoSMIM’s margin over these two methods is consistent with the idea that the gene–gene covariance of a niche carries domain information that per-gene statistics do not record.

### 3.2. Domain-specific gene–gene correlations obtained from the CoSMIM features

Here we search for gene pairs whose co-variation differs in strength or sign between spatial domains. Co-variation involves two genes at once, so a feature built on mean expression does not record it directly. We also measure how closely the correlations in the domains predicted by CoSMIM match those in the annotated domains. CoSMIM keeps a gene–gene covariance matrix for each cell, and each entry corresponds to one gene pair. Averaging each cell’s niche covariance **Σ**_*i*_ within a spatial domain and converting to a correlation matrix gives the strength of co-variation of every gene pair within that domain. Differential co-expression analysis is established for bulk and spatial data (Acharyya et al., 2022; Fukushima, 2013). In CoSMIM, the domain-level correlations are computed from the same features that are used for clustering, so no separate model needs to be fitted. We use dataset1, which has five sections and 64 selected genes (2016 gene pairs), to keep gene pair correlation results small enough to report (Fig. 3).

**Fig. 3:**
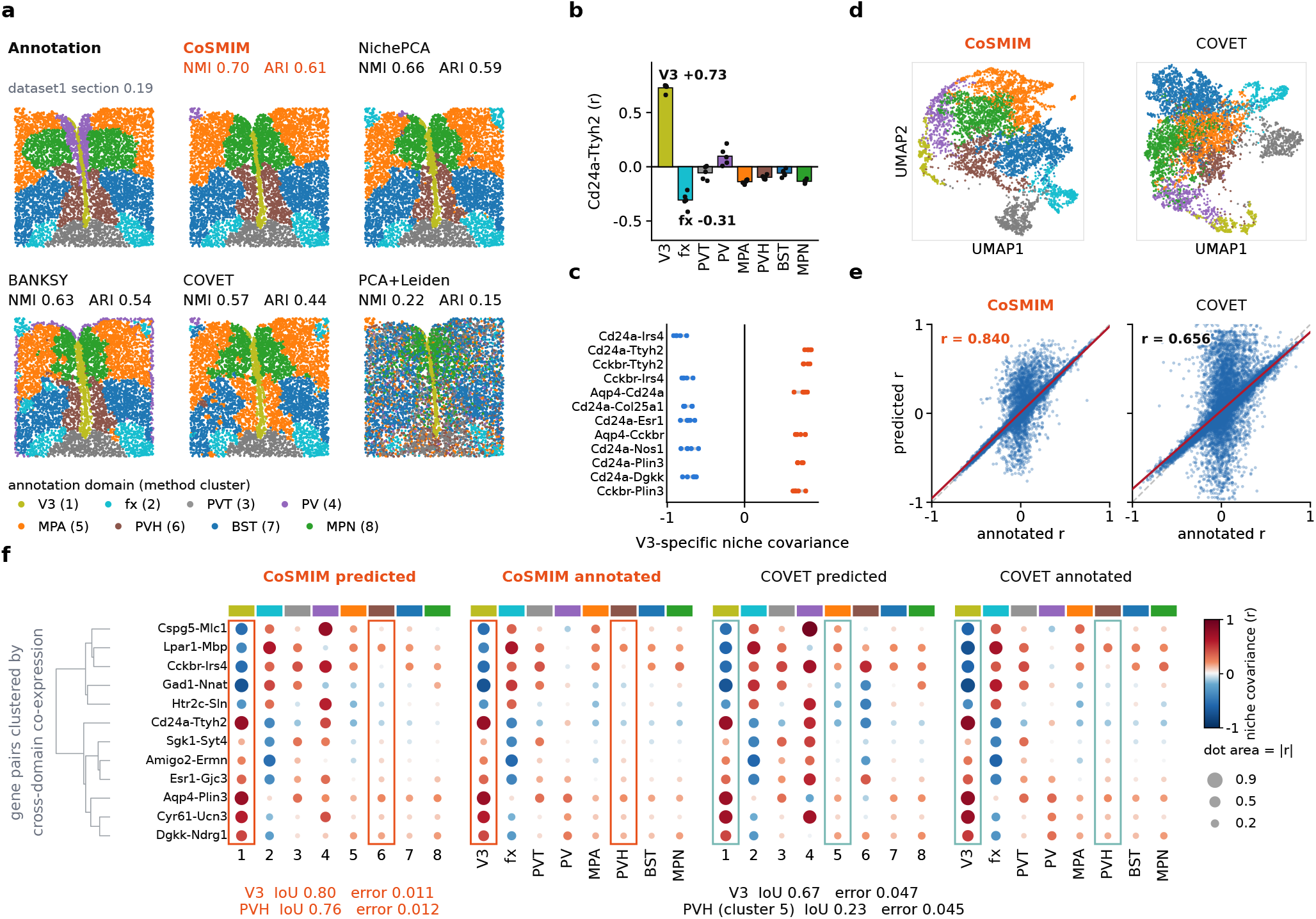
Gene–gene correlations on dataset1 (MERFISH mouse hypothalamus), section 0.19. **(a)** The annotation and the clusters of the five methods, with NMI and ARI. Each cluster has the colour of the annotated domain matched to it one-to-one. **(b)** Correlation of *Cd24a*–*Ttyh2* in each annotated domain. Bars show the mean over the five sections and points show single sections. **(c)** The twelve pairs with the largest absolute V3 contrast among the significant pairs tested on all five sections (§3.2). **(d)** UMAP of the CoSMIM and COVET features, coloured by the annotation. **(e)** Correlations of all 2016 gene pairs in each predicted domain against the matched annotated domain. The red line is the least-squares fit. **(f)** The twelve pairs with the largest range across domains in the CoSMIM annotated block, from the predicted and the annotated partition of each method. Boxes mark V3 and PVH. Below each predicted block are the IoU of the cluster that best overlaps each boxed domain and its mean absolute difference from that domain over all 2016 pairs.

We first compare the partitions of the five methods on one section (Fig. 3a), because the matrix-level comparison below (Fig. 3e,f) is computed on the predicted partitions. All five methods are run with a fixed protocol of §2.5 and each method’s clusters are relabelled to the annotated domain they best match in Fig. 3a. On this dataset1 section 0.19, CoSMIM ranks first at NMI 0.70 and ARI 0.61, ahead of all competitor methods. In the UMAP of the CoSMIM and COVET features coloured by the annotation (Fig. 3d), CoSMIM separates the eight domains better than COVET which overlaps several annotated domains.

A per-domain correlation matrix cannot be compared directly across sections, because a large part of the correlation structure is shared by all domains of the same section. We therefore use a within-section contrast, defined as the correlation of a gene pair in a domain minus the median of its correlations in the other domains of the same section. We score every gene pair this way on every section. A pair is tested on the sections that selected both of its genes. For each of the 1854 pairs tested on at least three sections, we use a one-sample *t*-test across those sections to test whether the mean contrast differs from zero. Pairs with a Benjamini–Hochberg adjusted *q* < 0.05 are called significant.

We applied this test to four annotated domains (V3, fx, PV and PVH), and two of them have gene pairs with a significant contrast.

Of the 1854 pairs tested, 946 have a significant contrast in V3, and 402 of these are tested on all five sections. The corresponding numbers are 892 (452) for fx, 163 (90) for PV and 5 (4) for PVH. The pairs with the largest absolute V3 contrast also vary little across sections (Fig. 3c). The top two pairs in Fig. 3c are *Cd24a*– *Irs4*, with a V3 contrast of −0.85 ± 0.06 (mean ± s.d. over the five sections, *q* = 0.0008), and *Cd24a*–*Ttyh2*, with +0.83 ± 0.03 (*q* = 0.0005). For both pairs, the correlation in fx has the opposite sign to that in V3 on all five sections. *Cd24a*–*Irs4* has a mean correlation of −0.62 in V3 and +0.40 in fx, and *Cd24a*–*Ttyh2* has +0.73 in V3 and −0.31 in fx (Fig. 3b). Both fx contrasts are also significant (*q* ≤ 0.008). The correlation of the same gene pair can therefore differ in sign between two domains. A first-order feature does not contain this quantity directly. However, the covariance is centred on the global mean, part of each correlation can come from the shift of the domain mean expression.

Eight of the twelve pairs in Fig. 3c contain *Cd24a*, and the other four contain *Cckbr*. Of the 96 genes selected on at least one section, *Cd24a* has the largest increase in mean expression in V3 over the other domains, and *Cckbr* has the fourth largest. *Cd24a* encodes CD24, which is present on the apical surface of most ependymal cells lining V3 (Mirzadeh et al., 2017). *Cckbr* encodes the cholecystokinin B receptor. *Irs4* encodes insulin receptor substrate 4. Neurons in and around the PVH that express *Irs4* are required for normal energy balance (Sutton Hickey et al., 2020). *Ttyh2* encodes a member of the tweety family of membrane proteins. The sign change of *Cd24a*–*Irs4* and *Cd24a*– *Ttyh2* between V3 and fx is consistent with the different cell compositions of the ependymal lining and the fornix fibre tract. It does not by itself show that the two genes of either pair regulate each other.

We next compare whole correlation matrices instead of single gene pairs using COVET. Like CoSMIM, COVET clusters PCA-reduced neighbourhood covariance features with Leiden but uses a single neighbourhood size (*k* = 8). The two methods also differ in gene selection, shrinkage and clustering parameters. We test whether the correlation matrices of the predicted domains agree with those of the annotated domains (Fig. 3e,f and Supplementary Fig. S3).

CoSMIM reproduces the eight annotated domains better than COVET, which splits some nuclei and merges others (Fig. 3a). Over the 64 genes shared by the two methods, the concordance is *r* = 0.840 for CoSMIM and *r* = 0.656 for COVET (Fig. 3e). Both differences are larger on dataset2. CoSMIM (NMI 0.71) recovers most of the cortical layers, the white matter and the hippocampus, whereas COVET assigns most of the cortex to a single cluster at NMI 0.46. Over all 496 pairs of the 32 selected genes, the concordance is *r* = 0.783 for CoSMIM and *r* = 0.264 for COVET (Supplementary Fig. S3).

Fig. 3f compares the two methods on two domains, V3 and PVH. For each method and domain, we use the cluster with the highest intersection over union (IoU) with the domain. CoSMIM recovers V3 at an IoU of 0.80. Over all 2016 gene pairs, the mean absolute difference between the correlations of this cluster and those of the annotated V3 is 0.011. The COVET cluster matched to V3 has an IoU of 0.67 and a mean absolute difference of 0.047. CoSMIM recovers PVH at an IoU of 0.76, with a difference of 0.012. COVET has no cluster of its own for PVH on this section. The correlations of the domains that CoSMIM predicts are therefore close to those of the annotated domains. This suggests that domain-level gene–gene correlations can be estimated from CoSMIM domains on sections without an annotation.

### 3.3. A spatial domain resolved by CoSMIM, but not by mean expression or single-scale covariance

Organs in the early mouse embryo can be small and consist of several cell lineages. Averaging expression over such a mixture may give a niche profile that differs little from those of other tissues. These organs are therefore a stringent test of features built on mean expression. To assess method performance on the mouse embryo Stereo-seq dataset 6, we evaluated whether derived clusters accurately captured each annotated organ across tissue sections. Alignment precision was quantified using the IoU between an organ annotation and its best-aligned cluster, classifying any target with an IoU score below 0.3 as unresolved. For the brain on section <monospace>E10.5 E2S1</monospace>, all methods except PCA+Leiden resolved it, and CoSMIM had the highest IoU (Fig. 4a). However, for the liver on section <monospace>E10.5 E1S1</monospace>, CoSMIM was the only method that resolved it (Fig. 4b). CoSMIM reached an IoU of 0.79, whereas all other methods scored between 0.03 and 0.10. This liver region was the smallest annotated domain on the section <monospace>E10.5 E1S1</monospace>, comprising only 195 of the 18,404 total cells. A similar result of liver identification was observed on E9.5 sections.

**Fig. 4:**
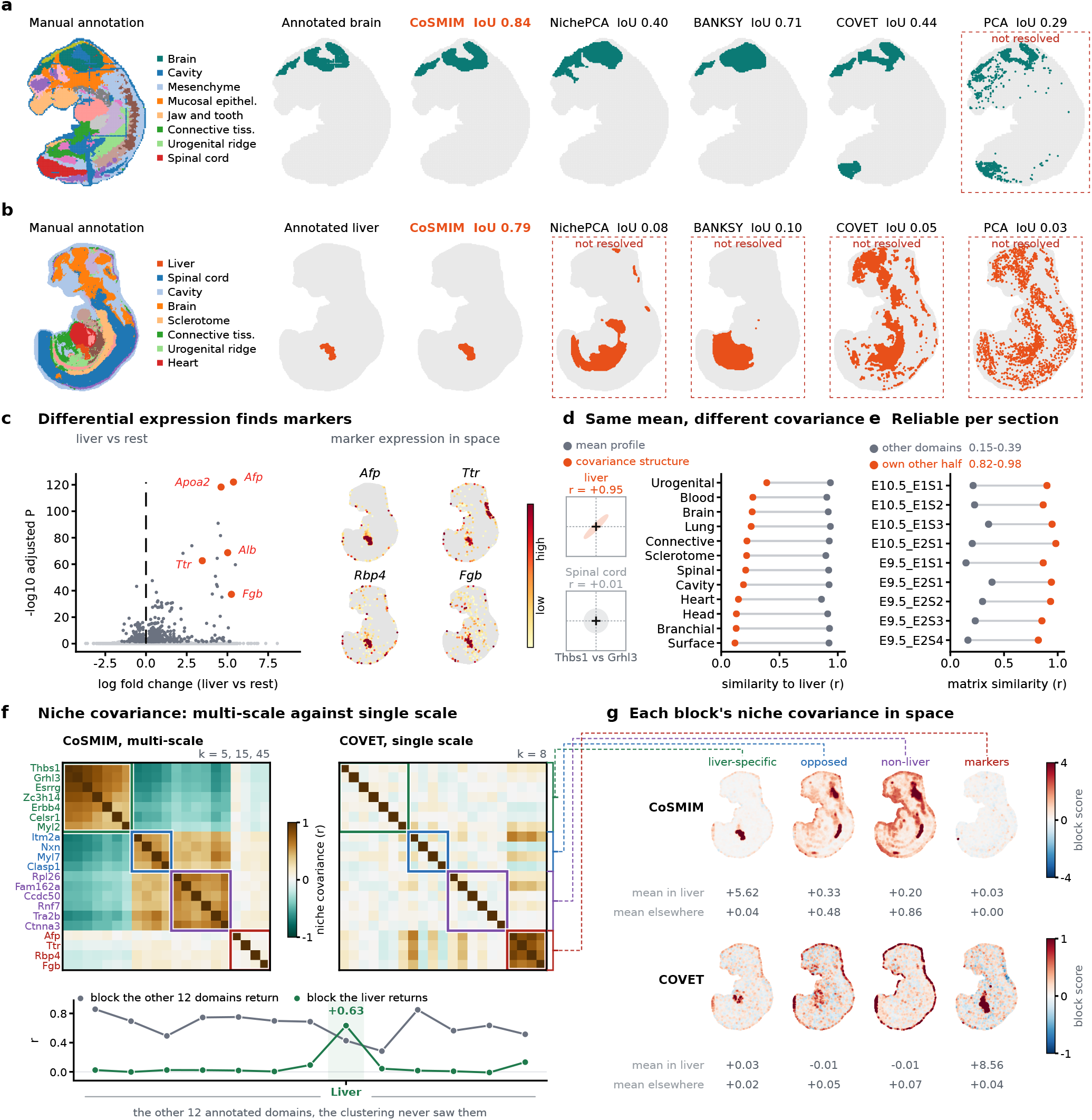
dataset6 (Stereo-seq mouse embryo). **(a, b)** The brain on E10.5 E2S1 and the liver on E10.5 E1S1, with the best-matching cluster of each method and its IoU. Red dashed boxes mark IoU*<* 0.3. **(c)** The liver against all other domains (Wilcoxon test), and four hepatoblast markers in space. **(d)** Left, one gene pair in two domains, standardised so that only the correlation differs. Right, the similarity of each domain to the liver by mean profile (grey) and by covariance (orange). **(e)** Correlation between the liver matrices of two random halves of the liver cells (orange) and median correlation with the other domains (grey), on nine sections. **(f)** Upper, the liver niche covariance over 21 genes for CoSMIM and COVET. The green, blue and purple boxes are the blocks of Supplementary Fig. S4, and the red box marks the markers of (c). Lower, the mean within-block correlation of the liver block (green) and the shared set (grey) in each domain. **(g)** The block scores in space. Panels (c), (d), (f) and (g) show E10.5 E1S1.

We then applied a differential expression test to the same section. The differential expression test comparing the liver against the rest of the tissue identified canonical abundance-based marker genes, including *Afp, Ttr, Rbp4* and *Fgb*, colocalizing to the same spatial region (Fig. 4c). Because these genes represent established markers of the hepatoblast and early fetal-liver programme (Su et al., 2017), the domain CoSMIM alone recovers is independently confirmed to be the liver. However, a differential expression analysis identifies highly expressed genes in a region rather than delineating cell functional spatial domains.

The CoSMIM identification of the liver relies on second-order statistics rather than only first-order statistics. To evaluate separability, we compared the liver niche with each remaining annotated domain on the same section by Pearson correlation *r*, where a higher *r* indicates lower separability. The median *r* is 0.92 for the mean expression profiles and 0.21 for the niche covariance matrices (Fig. 4d). The mean expression of each gene is not centred before this *r* is computed, so the mean-profile *r* largely reflects the baseline expression level of each gene. This clear separation is consistently observed across all nine sections, even when clustering fails to recover the organ, indicating that organ recovery is constrained by the partitioning algorithm rather than the underlying CoSMIM feature representation. The per-domain estimate is highly stable: across all nine sections, splitting the liver cells in half and estimating the niche CoSMIM independently for each subset gives high internal consistency at *r* = 0.82–0.98, compared to low correlations 0.15–0.39 between the liver CoSMIM matrix and the other domains on the same section (Fig. 4e).

Clustering the liver’s own CoSMIM matrix correlation (Supplementary Fig. S4) returns a tight gene block that has the highest mean within-block correlation *r* at 0.633 (Fig. 4f green box). However, only evaluating this block in isolation is insufficient to demonstrate biological specificity. Applying the identical clustering procedure to all thirteen annotated domains returns a tight purple block, with mean within-block correlations from +0.60 to +0.86. Because the liver’s correlation ranks eleventh of thirteen (Fig. 4f green line x-axis liver point), high within-block correlation reflects an inherent artifact of the optimization algorithm rather than organ-specific evidence. Instead, specificity is evidenced by gene identity. Twelve genes appear in the tightest block of at least eight of the other twelve domains. We call them the shared set (Fig. 4f, grey line). The tightest blocks of eleven of these twelve domains overlap the shared set. This liver-specific set showed an average correlation of +0.633 inside the liver, but ranged from −0.009 to +0.133 across the remaining twelve domains (Fig. 4f green line). Because the clustering was performed solely on the liver, the reduction of cross-domain correlation to near zero provides independent out-of-sample validation that this module is confined to the liver.

The block of the four canonical marker genes of Fig. 4c in CoSMIM is distinct from the differential-expression program (Fig. 4f red box). The red block exhibits a within-block correlation of +0.046 substantially below those of the thirteen domain-derived blocks—indicating minimal spatial co-variation. Therefore, genes that are highly expressed in the liver do not necessarily co-vary strongly in the liver niche. The E10.5 liver bud is a heterogeneous aggregate comprising hepatoblasts delaminating from the foregut endoderm into the septum transversum mesenchyme, together with endothelial cells and colonising haematopoietic cells (Gordillo et al., 2015). Consequently, niche CoSMIM is dominated by spatial co-occurrence of these cell lineages rather than by the single-cell marker elevation. In summary, the liver-restricted block indicates a covariation structure that is spatially specific to the liver on this tissue section.

We then mapped the score of each block onto the section (Fig. 4g). A cell’s block score corresponds to its average niche covariance entry. Based on this score, the CoSMIM liver-specific block is selectively elevated inside the liver and absent outside, whereas the consensus block (non-liver) shared by the other twelve domains exhibits the opposite pattern, peaking in the body cavity, vasculature, and sclerotome. Under single-scale COVET, on the same cells and genes, that liver-specific block is as flat inside the liver as outside and does not separate the organ. Instead, the marker block becomes its strongest structure. Both methods centre each neighbourhood on the global mean, so genes jointly elevated within a domain naturally produce positive cross-terms (§2.1). Single-scale COVET seems to re-encode the marker gene first-order signals. Our CoSMIM retains a spatial covariation feature that remains consistent across multiple scales. The liver bud is a small domain made up of several cell lineages. Its niche covariance differs from those of the other domains much more than its mean expression profile does (Fig. 4d).

## 4. Discussion and conclusion

CoSMIM summarises a cell’s microenvironment by the gene–gene covariance of its neighbourhood at several scales, taking the matrix square root of the fused covariance as the feature. Standard PCA and Leiden clustering of this feature identify spatial domains without model training and attain the best overall ranking on the six public benchmarks from five platforms and the highest median NMI on the Xenium mouse brain dataset. On one section of the Stereo-seq mouse embryo dataset (E10.5 E1S1), CoSMIM is the only one of five methods that resolves the liver. In addition, because the feature is built from covariance matrices, domain-specific gene–gene correlations can be obtained from these matrices and tested across sections. The covariance of a neighbourhood can record which cell populations share it and which genes distinguish them. The mean reduces this mixture to a single profile. Our two examples fit this view. In the hypothalamus the top gene pairs have correlations of opposite sign in the third ventricle and the fornix.

This sign change is in line with the different cell compositions of the ependymal lining and the fibre tract. In the embryo the liver bud is a small domain that contains hepatoblasts, endothelial cells and haematopoietic cells (Gordillo et al., 2015). It is separated from the other domains by its niche covariance. The per-domain matrices can therefore suggest gene pairs for each domain. Pairs that recur across sections are candidates for experimental follow-up.

The scale set K is still chosen by hand and the three values of *k* rest on one ablation, so another platform or tissue will likely need retuning. Future work could set K from the spatial resolution and cell density of the data. Another direction is to cluster on the symmetric positive definite cone itself, along its geodesics rather than Euclidean lines in square-root coordinates.

## Acknowledgements

The computational resources of the supercomputer were provided by the Human Genome Center, the Institute of Medical Science, The University of Tokyo. Bairui Du is supported as a research assistant by the World-leading Innovative Graduate Study Program on Global Leadership for Social Design and Management (WINGS-GSDM), The University of Tokyo.

## Author contributions statement

Bairui Du conceived the method, implemented the software, performed the analyses and wrote the manuscript. Hisanori Kiryu supervised the project, acquired the funding and revised the manuscript. Both authors read and approved the final manuscript.

## Data availability

All data is publicly available and was processed as described in Section 2.

