## Supplementary Material: Supplementary Notes S1-S5 and Supplementary Figures S1-S8 for "Multi-scale microenvironment covariance for unsupervised spatial domain identification"

Bairui Du and Hisanori Kiryu

The Supplementary Material has two parts, Supplementary Notes S1–S5 and Supplementary Figures S1–S8. The tables are inside the Notes. Supplementary Data S1 is a separate workbook. It lists every section with its source, number of cells, genes and annotated domains, gives the metrics and ranks of every method on every section and dataset, and records every setting of CoSMIM and of the competitors. It also gives the scores of the competitors at each $k$ from which their $k$ was chosen.

#### Supplementary Notes

### Supplementary Note S1: The CoSMIM feature

##### Notation

$N$ is the number of cells in a section and $g$ is the number of genes used for the covariance. $\mathbf{x}_r \in \mathbb{R}^g$ is the log-expression of cell $r$ after preprocessing, and $\boldsymbol{\mu} = \frac{1}{N}\sum_{r=1}^{N}\mathbf{x}_r$ is the global mean over the section. The scales are $\mathcal{K} = \{k_1 < \cdots < k_m\}$. Every benchmark run uses $\mathcal{K} = \{5, 15, 45\}$. $\mathcal{N}^{(j)}(i)$ is the set of cell $i$ and its $k_j$ nearest neighbours in space, so it has $k_j + 1$ cells. A matrix carries the scale index $j$ as a superscript in parentheses. The scalars $k_j$ and $w_j$ carry it as a subscript. $w_j$ are the scale weights, $\alpha = 0.1$ is the shrinkage strength, $\mathrm{vech}(\cdot)$ takes the lower triangle of a symmetric matrix together with its diagonal, and $\|\mathbf{M}\|_F = \mathrm{tr}(\mathbf{M}^\top\mathbf{M})^{1/2}$ is the Frobenius norm.

##### Construction

The covariance of cell $i$ at scale $k_j$ is centred on the global mean $\boldsymbol{\mu}$:

$$\mathbf{C}_i^{(j)} = \frac{1}{k_j}\sum_{r\in\mathcal{N}^{(j)}(i)} (\mathbf{x}_r - \boldsymbol{\mu})(\mathbf{x}_r - \boldsymbol{\mu})^\top + \varepsilon\,\mathbf{I}. \tag{S1}$$

The divisor $k_j$ is the number of cells minus one, as in the COVET implementation (Haviv et al., 2025). The ridge $\varepsilon\mathbf{I}$ keeps the matrix invertible when $k_j < g$. $\varepsilon$ is $10^{-5}$ times the mean of all entries of the scale-$k_j$ matrices before the ridge is added, taken over the cells of one computing batch (at most 1000 cells). The scales are then fused, shrunk and mapped by the matrix square root:

$$\boldsymbol{\Sigma}_i^w = \sum_{j=1}^{m} w_j\,\mathbf{C}_i^{(j)}, \qquad w_j \ge 0,\ \textstyle\sum_j w_j = 1, \tag{S2}$$

$$\boldsymbol{\Sigma}_i = (1-\alpha)\,\boldsymbol{\Sigma}_i^w + \alpha\,\frac{\mathrm{tr}(\boldsymbol{\Sigma}_i^w)}{g}\,\mathbf{I}, \tag{S3}$$

$$\mathbf{S}_i = \boldsymbol{\Sigma}_i^{1/2} = \mathbf{U}\,\boldsymbol{\Lambda}^{1/2}\,\mathbf{U}^\top, \qquad \boldsymbol{\Sigma}_i = \mathbf{U}\boldsymbol{\Lambda}\mathbf{U}^\top. \tag{S4}$$

$\boldsymbol{\Sigma}_i$ is the CoSMIM matrix of cell $i$, and the feature passed to PCA is $\mathrm{vech}(\mathbf{S}_i)$. The shrinkage of Eq. (S3) uses the scaled identity target of Ledoit and Wolf (2004). The strength $\alpha = 0.1$ is the same on all datasets and was not tuned for any one dataset.

#### The first-order term

Let $\bar{\mathbf{x}}_i^{(j)}$ be the mean expression of $\mathcal{N}^{(j)}(i)$ and $\mathbf{d}_i^{(j)} = \bar{\mathbf{x}}_i^{(j)} - \boldsymbol{\mu}$. Because the cross terms sum to zero, the first term of Eq. (S1) splits into two parts:

$$\begin{aligned}\frac{1}{k_j} \sum_{r\in\mathcal{N}^{(j)}(i)} &(\mathbf{x}_r - \boldsymbol{\mu})(\mathbf{x}_r - \boldsymbol{\mu})^\top \\ &= \frac{1}{k_j} \sum_{r\in\mathcal{N}^{(j)}(i)} (\mathbf{x}_r - \bar{\mathbf{x}}_i^{(j)})(\mathbf{x}_r - \bar{\mathbf{x}}_i^{(j)})^\top + \frac{k_j+1}{k_j}\,\mathbf{d}_i^{(j)}\mathbf{d}_i^{(j)\top}. \end{aligned} \tag{S5}$$

The first part is the covariance around the neighbourhood mean, a second-order statistic. The second part depends only on the neighbourhood mean, so it is a first-order term. This is the first-order term mentioned in the Introduction (§1). Centring on $\boldsymbol{\mu}$ instead of on the neighbourhood mean gives the matrices of all niches one common origin, as in COVET (Haviv et al., 2025).

#### Scale weights

One weight vector is learned for each section and then used for every cell of that section. Let $\mathcal{R}$ be a fixed random subset of at most 1000 cells of the section and $\bar{\mathbf{C}}^{(j)} = \frac{1}{|\mathcal{R}|}\sum_{i\in\mathcal{R}} \mathbf{C}_i^{(j)}$. The weights minimise

$$\mathcal{L}(\mathbf{w}) = -\log\det\Big(\sum_{j=1}^{m} w_j\,\bar{\mathbf{C}}^{(j)}\Big) + \lambda \sum_{j=1}^{m} w_j \log(w_j + \eta) \tag{S6}$$

over $w_j \geq 0$, $\sum_j w_j = 1$, where $\eta = 10^{-10}$ keeps the logarithm finite. The problem is solved by the SLSQP solver of SciPy (Virtanen et al., 2020) with the approximate gradient $\partial\mathcal{L}/\partial w_j \approx -\operatorname{tr}\big((\bar{\boldsymbol{\Sigma}}^w)^{-1}\bar{\mathbf{C}}^{(j)}\big) + \lambda\big(\log(w_j+\eta)+1\big)$, where $\bar{\boldsymbol{\Sigma}}^w = \sum_j w_j \bar{\mathbf{C}}^{(j)}$. The first term is smaller when $\bar{\boldsymbol{\Sigma}}^w$ has a larger determinant. The second term is smallest at equal weights, $w_j = 1/m$. A larger $\lambda$ moves the solution towards equal weights. The value of $\lambda$ on each dataset is in Supplementary Data S1.

#### Square-root distance and the Bures–Wasserstein distance

The matrices $\boldsymbol{\Sigma}_i$ are symmetric positive definite. The set of such matrices is a cone and not a linear space (Pennec et al., 2006). A covariance matrix can be pictured as an ellipsoid. Its axes show which genes vary together and by how much. A natural distance between two covariance matrices $\mathbf{A}$ and $\mathbf{B}$ is the Bures–Wasserstein distance, the optimal-transport distance between the zero-mean Gaussian distributions with these covariances (Bhatia et al., 2019):

$$d^2_{\mathrm{BW}}(\mathbf{A},\mathbf{B}) = \operatorname{tr}(\mathbf{A}) + \operatorname{tr}(\mathbf{B}) - 2\operatorname{tr}\big[(\mathbf{A}^{1/2}\mathbf{B}\mathbf{A}^{1/2})^{1/2}\big]. \tag{S7}$$

As in COVET (Haviv et al., 2025), CoSMIM approximates this distance by the Euclidean distance between the matrix square roots. Let $\mathbf{M} = \mathbf{A}^{1/2} - \mathbf{B}^{1/2}$, which is symmetric. Then

$$\|\mathbf{A}^{1/2} - \mathbf{B}^{1/2}\|_F^2 = \operatorname{tr}(\mathbf{M}^2) = \operatorname{tr}(\mathbf{A}) + \operatorname{tr}(\mathbf{B}) - 2\operatorname{tr}(\mathbf{A}^{1/2}\mathbf{B}^{1/2}), \tag{S8}$$

using $\mathbf{A}^{1/2}\mathbf{A}^{1/2} = \mathbf{A}$ and $\operatorname{tr}(\mathbf{XY}) = \operatorname{tr}(\mathbf{YX})$. Eqs. (S7) and (S8) differ only in the cross term.

**Equal when the matrices commute.** If $\mathbf{AB} = \mathbf{BA}$, both matrices are diagonal in one orthonormal basis, with eigenvalues $a_l$ and $b_l$. Both cross terms are then $\sum_l \sqrt{a_l b_l}$, so $\|\mathbf{A}^{1/2} - \mathbf{B}^{1/2}\|_F = d_{\mathrm{BW}}(\mathbf{A},\mathbf{B})$.

**Never smaller in general.** Let $\mathbf{P} = \mathbf{A}^{1/2}\mathbf{B}^{1/2}$. Then $\mathbf{PP}^\top = \mathbf{A}^{1/2}\mathbf{B}\mathbf{A}^{1/2}$, so the cross term of Eq. (S7) is $\operatorname{tr}[(\mathbf{PP}^\top)^{1/2}]$, the nuclear norm $\|\mathbf{P}\|_*$. The trace of a matrix is not larger than its nuclear norm, so $\operatorname{tr}(\mathbf{A}^{1/2}\mathbf{B}^{1/2}) \leq \operatorname{tr}[(\mathbf{A}^{1/2}\mathbf{B}\mathbf{A}^{1/2})^{1/2}]$ and

$$\|\mathbf{A}^{1/2} - \mathbf{B}^{1/2}\|_F \;\geq\; d_{\mathrm{BW}}(\mathbf{A},\mathbf{B}), \tag{S9}$$

Table S1: **dataset4 against the two annotations.** NMI and ARI of the five methods on section BY3. The benchmark uses the SDMBench annotation (ground_truth). The competitors are at the global $k$ of the benchmark. For BANKSY, the two annotations score two runs at $k = 20$. For the other methods, they score the same clusters. The largest value of each column is in bold.

| | SDMBench annotation | | smoothed_manual | |
|---|---|---|---|---|
| Method | NMI | ARI | NMI | ARI |
| CoSMIM | **0.731** | **0.639** | 0.775 | **0.763** |
| NichePCA ($k = 15$) | 0.689 | 0.574 | **0.807** | 0.743 |
| BANKSY ($k = 20$) | 0.637 | 0.502 | 0.751 | 0.657 |
| COVET ($k = 8$) | 0.617 | 0.456 | 0.689 | 0.550 |
| PCA+Leiden | 0.252 | 0.213 | 0.290 | 0.248 |

with equality when $\mathbf{A}$ and $\mathbf{B}$ commute.

The map $\mathbf{A} \mapsto \mathbf{A}^{1/2}$ is one-to-one on symmetric positive definite matrices, so the square-root distance is a metric and is zero only when the two covariances are equal. Because $\text{vech}(\mathbf{S}_i)$ is an ordinary real vector, PCA, the $k$NN graph and Leiden (Traag et al., 2019) can be applied to it without change. The implementation does not multiply the off-diagonal entries of vech by $\sqrt{2}$. The Euclidean distance between two feature vectors therefore gives each off-diagonal entry half of its weight in the squared Frobenius norm. It is still a metric. This paper does not cluster with the Bures–Wasserstein distance itself or along the geodesics of the cone.

### Supplementary Note S2: Datasets and annotations

Supplementary Data S1 lists the seven datasets section by section. For dataset1 to dataset6 we use the annotations distributed with SDMBench (Yuan et al., 2024), which NichePCA (Schaub et al., 2025) also uses. The eight domains of dataset1, the seven domains of dataset4 and the layers of dataset5 were annotated by Li and Zhou (2022). The annotations of dataset2, dataset3 and dataset6 come from the original studies (Codeluppi et al., 2018; Long et al., 2023; Chen et al., 2022). For dataset7 we use the region annotation of Bhuva et al. (2024), who registered each section to the Allen mouse brain atlas (Wang et al., 2020) (140 to 144 regions). For dataset7 we score the left hemisphere and set the target number of clusters to 60 (Section 2.5). The observations of dataset6 are bins of the Stereo-seq grid, which the text calls cells.

Singhal et al. (2024) score dataset4 against a second annotation, smoothed_manual, which they derived from the cell types of Wang et al. (2018) by label smoothing. Supplementary Table S1 therefore also gives the comparison of CoSMIM and the competitors on smoothed_manual. On smoothed_manual, NichePCA has the highest NMI and CoSMIM has the highest ARI.

### Supplementary Note S3: Ablations of the design

Three ablations test the choices of §2.1: the number of scales, the sizes of the scales and the map from a covariance matrix to a Euclidean feature. Each ablation uses its own settings. Only the three-scale entries of dataset1 and dataset2 in Supplementary Table S2 are benchmark runs.

**Number of scales (Supplementary Table S2).** For dataset1 and dataset2, each entry is the best median NMI over a grid of the number of principal components and the number of neighbours of the clustering graph (Supplementary Fig. S1). On these two datasets, more than three scales give no gain larger than 0.003. For dataset7, each entry is the best of the available runs, and NMI increases up to five scales. The benchmark uses three scales on all seven datasets, so that one scale set serves every dataset. More scales also take longer to compute, because the covariance step costs $O(N\, m\, k_m\, g^2)$ (caption of Supplementary Fig. S8).

Table S2: **Number of scales** (median NMI). dataset1 and dataset2: the best value over the grid of Supplementary Fig. S1, with learned weights. dataset7: the best of the available runs at 20 or 30 principal components and 5 to 100 graph neighbours, with uniform weights and scored on whole sections, so this column is not on the same basis as the dataset7 benchmark (learned weights, left hemisphere).

| Scales | dataset1 | dataset2 | dataset7 |
|---|---|---|---|
| three, $[5, 15, 45]$ | **0.68** | **0.71** | 0.64 |
| four, $[5, 15, 45, 135]$ | 0.67 | 0.69 | 0.67 |
| five, $[5, 15, 45, 135, 405]$ | 0.65 | **0.71** | **0.68** |

Table S3: **Sizes of the scales** (three scales, median NMI). All sets use 30 principal components and 50 (dataset1) or 75 (dataset2) neighbours of the clustering graph.

| Scale set | dataset1 | dataset2 |
|---|---|---|
| $[5, 15, 45]$ (used in this paper) | 0.587 | 0.639 |
| $[3, 9, 27]$ | 0.569 | 0.596 |
| $[8, 24, 72]$ | 0.521 | 0.626 |
| $[5, 17, 53]$ (coprime) | **0.598** | **0.648** |

Table S4: **Map from a covariance matrix to a Euclidean feature** (median NMI, the same settings for the three maps on each dataset). The log-Euclidean map is that of Arsigny et al. (2006). dataset4 is scored against the annotation used in the benchmark (Supplementary Note S2). On dataset4 the three maps use the scales $\{8, 15, 50\}$, $\lambda = 1$, 20 principal components and 50 graph neighbours, not the benchmark settings, so the square-root value differs from Supplementary Table S1.

| Map | dataset1 | dataset4 |
|---|---|---|
| matrix square root $\mathbf{\Sigma}^{1/2}$ (CoSMIM) | **0.659** | **0.716** |
| matrix logarithm $\log \mathbf{\Sigma}$ (log-Euclidean) | 0.562 | 0.609 |
| raw covariance entries | 0.434 | 0.671 |

**Sizes of the scales (Supplementary Table S3).** The number of scales is three and the downstream settings are fixed. On both datasets the two best sets are $[5, 15, 45]$ and the coprime set $[5, 17, 53]$. The coprime set is higher by 0.011 on dataset1 and by 0.009 on dataset2. We keep $[5, 15, 45]$.

**Map to a Euclidean feature (Supplementary Table S4).** At the same settings, the matrix square root gives the highest NMI on both datasets tested. The gap to the raw covariance entries is larger on dataset1 (0.659 against 0.434) than on dataset4 (0.716 against 0.671). A possible reason that the log-Euclidean map scores lower is that the logarithm enlarges the differences between small eigenvalues more than the square root does, and these eigenvalues are the least reliable in a covariance estimated from few cells.

#### Supplementary Note S4: Evaluation metrics

$N$ cells have spatial coordinates $\mathbf{s}_i$. $P$ is the predicted partition and $T$ is the annotation (Supplementary Note S2).

**Accuracy.** With the entropy $H(\cdot)$ and the mutual information $I(\cdot;\cdot)$,

$$\mathrm{NMI}(T,P) = \frac{2\,I(T;P)}{H(T)+H(P)}, \tag{S10}$$

$$\mathrm{HOM}(T,P) = 1 - \frac{H(T \mid P)}{H(T)}, \qquad \mathrm{COM}(T,P) = 1 - \frac{H(P \mid T)}{H(P)}. \tag{S11}$$

These three lie in $[0,1]$. HOM and COM are those of Rosenberg and Hirschberg (2007). With this normalisation, NMI equals their V-measure. Let $n_{uv} = |T_u \cap P_v|$, $a_u = \sum_v n_{uv}$ and $b_v = \sum_u n_{uv}$, and let $c_T = \sum_u \binom{a_u}{2}$, $c_P = \sum_v \binom{b_v}{2}$ and $c_N = \binom{N}{2}$. The adjusted Rand index (Hubert and Arabie, 1985) compares the two partitions over all pairs of cells and is corrected for chance:

$$\mathrm{ARI}(T,P) = \frac{\sum_{uv} \binom{n_{uv}}{2} - c_T c_P / c_N}{\frac{1}{2}(c_T + c_P) - c_T c_P / c_N}. \tag{S12}$$

ARI is 1 for identical partitions and close to 0 for a random partition. It lies in $[-1,1]$. The four accuracy metrics are computed with `scikit-learn` (Pedregosa et al., 2011).

**Continuity.** $\mathcal{N}_1(i)$ is the nearest cell to cell $i$ among the cells of its predicted cluster, and $\mathcal{M}(i)$ is the set of the 10 cells nearest to cell $i$.

$$\mathrm{CHAOS} = \frac{1}{N}\sum_{i=1}^{N} \left\| \mathbf{s}_i - \mathbf{s}_{\mathcal{N}_1(i)} \right\|, \tag{S13}$$

$$\mathrm{PAS} = \frac{1}{N}\sum_{i=1}^{N} \nVdash\Big[ \textstyle\sum_{l \in \mathcal{M}(i)} \nVdash[P_l \neq P_i] > 5 \Big]. \tag{S14}$$

CHAOS and PAS were introduced by Shang and Zhou (2022). Lower is better for both. For CHAOS the coordinates are first scaled to unit variance along each axis. ASW is the average silhouette width (Rousseeuw, 1987) of the predicted clusters, computed on the spatial coordinates. We report it rescaled to $[0,1]$ as $(\mathrm{ASW}+1)/2$. The published SDMBench values are rescaled in the same way before any ranking.

**Marker.** For one gene with log-normalised values $x_i$ and the weights $v_{il}$ of the spatial graph of the six nearest cells, with $W = \sum_{i \neq l} v_{il}$,

$$I = \frac{N}{W} \cdot \frac{\sum_{i \neq l} v_{il}(x_i - \bar{x})(x_l - \bar{x})}{\sum_i (x_i - \bar{x})^2}, \tag{S15}$$

$$C = \frac{N-1}{2W} \cdot \frac{\sum_{i \neq l} v_{il}(x_i - x_l)^2}{\sum_i (x_i - \bar{x})^2}. \tag{S16}$$

The genes are the top five marker genes of each predicted domain. Moran's $I$ (Moran, 1950) and Geary's $C$ (Geary, 1954) are the medians over these genes and are computed with `Squidpy` (Palla et al., 2022). Higher is better for Moran's $I$ and lower is better for Geary's $C$.

**Overall rank.** Within each metric the methods are ranked so that the largest rank is the best. SDMBench does not report ARI. The ranking with the eleven SDMBench methods therefore uses the other eight metrics, which weights accuracy, continuity and marker $3:3:2$. ARI is compared only among the five methods that we ran.

#### Supplementary Note S5: Statistical tests

The two tests of Section 2.5 compare the NMI of CoSMIM with that of NichePCA ($k = 15$), BANKSY ($k = 20$), COVET ($k = 8$, at most 64 genes) and PCA+Leiden on the 25 sections of the seven datasets. The unit of both tests is the section. dataset7 is scored on the left hemisphere (Section 2.5). We use the paired Wilcoxon signed-rank test (Wilcoxon, 1945), and the Friedman test (Friedman, 1937) with the Nemenyi post-hoc test (Demšar, 2006).

**Paired Wilcoxon signed-rank test.** CoSMIM has a higher NMI than NichePCA, BANKSY, COVET and PCA+Leiden on 22, 22, 24 and 24 of the 25 sections, respectively. The median paired difference in NMI (CoSMIM minus the competitor) is 0.030, 0.037, 0.090 and 0.238, in the same order. The two-sided test gives $P = 1.4 \times 10^{-4}$, $9.1 \times 10^{-4}$, $8.3 \times 10^{-7}$ and $1.2 \times 10^{-7}$. After a Bonferroni correction for the four comparisons (multiplying by four), the largest is $3.6 \times 10^{-3}$.

**Friedman test and Nemenyi post-hoc test.** Over the five methods and the 25 sections, the Friedman test gives $\chi^2 = 65.8$ and $P = 1.7 \times 10^{-13}$. The mean rank, with rank 1 for the highest NMI, is 1.32 for CoSMIM, 2.56 for BANKSY, 2.60 for NichePCA, 3.92 for COVET and 4.60 for PCA+Leiden. The Nemenyi test separates CoSMIM from NichePCA ($P = 0.034$), BANKSY ($P = 0.044$), COVET ($P = 6.1 \times 10^{-8}$) and PCA+Leiden ($P = 2.2 \times 10^{-12}$). The Nemenyi test controls the family-wise error rate over all pairs of methods, so it is stricter than the Wilcoxon test without correction.

**Per dataset.** CoSMIM has the highest median NMI of the five methods on all seven datasets (Supplementary Data S1). The comparison with COVET does not isolate the effect of the scales, because the two methods differ in more than the number of scales (Section 2.2).

#### Supplementary Figures

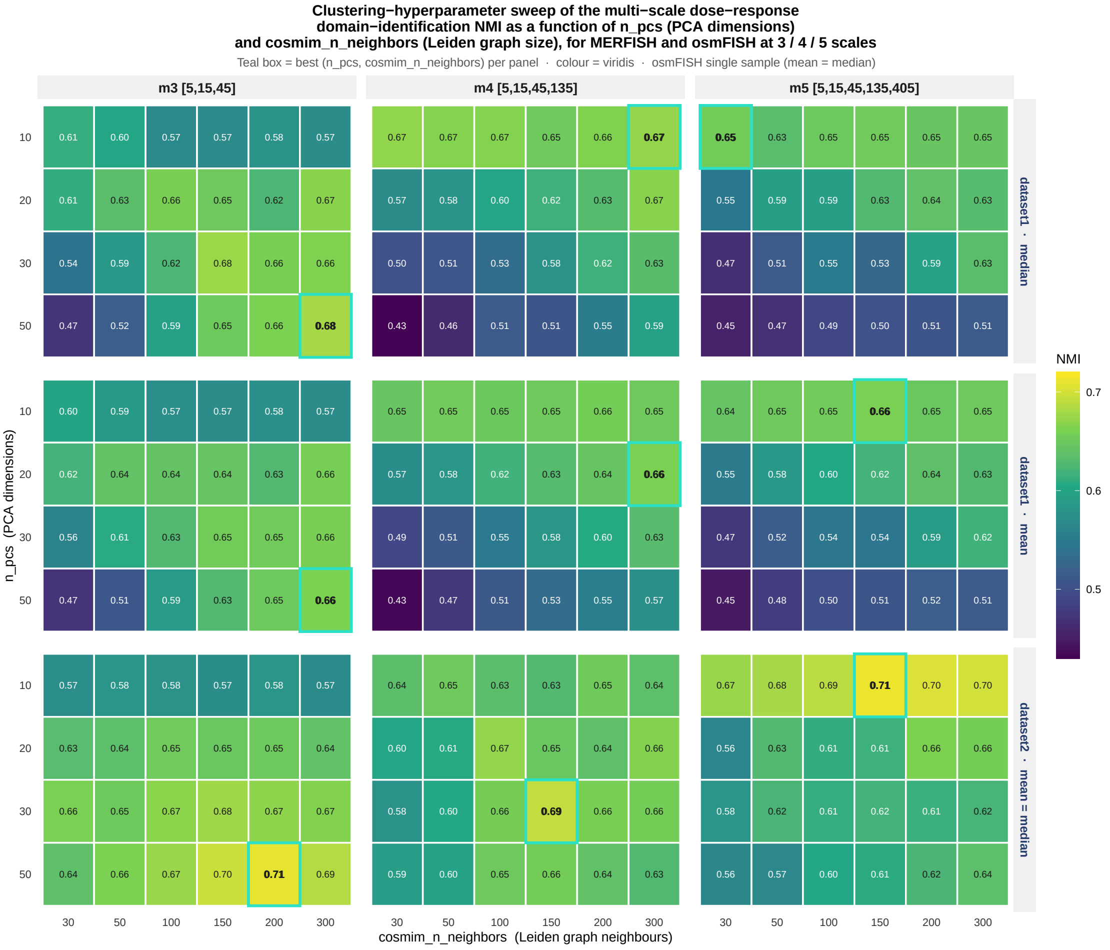


Figure S1: **Downstream settings for three, four and five scales.** NMI for each number of principal components (rows) and number of neighbours of the clustering graph (columns), with three (m3), four (m4) and five (m5) scales. The top row is the median over the five sections of dataset1, the middle row is the mean over the same sections and the bottom row is the one section of dataset2. The box marks the best cell of each panel. The boxed values of the top and bottom rows are the dataset1 and dataset2 entries of Supplementary Table S2.

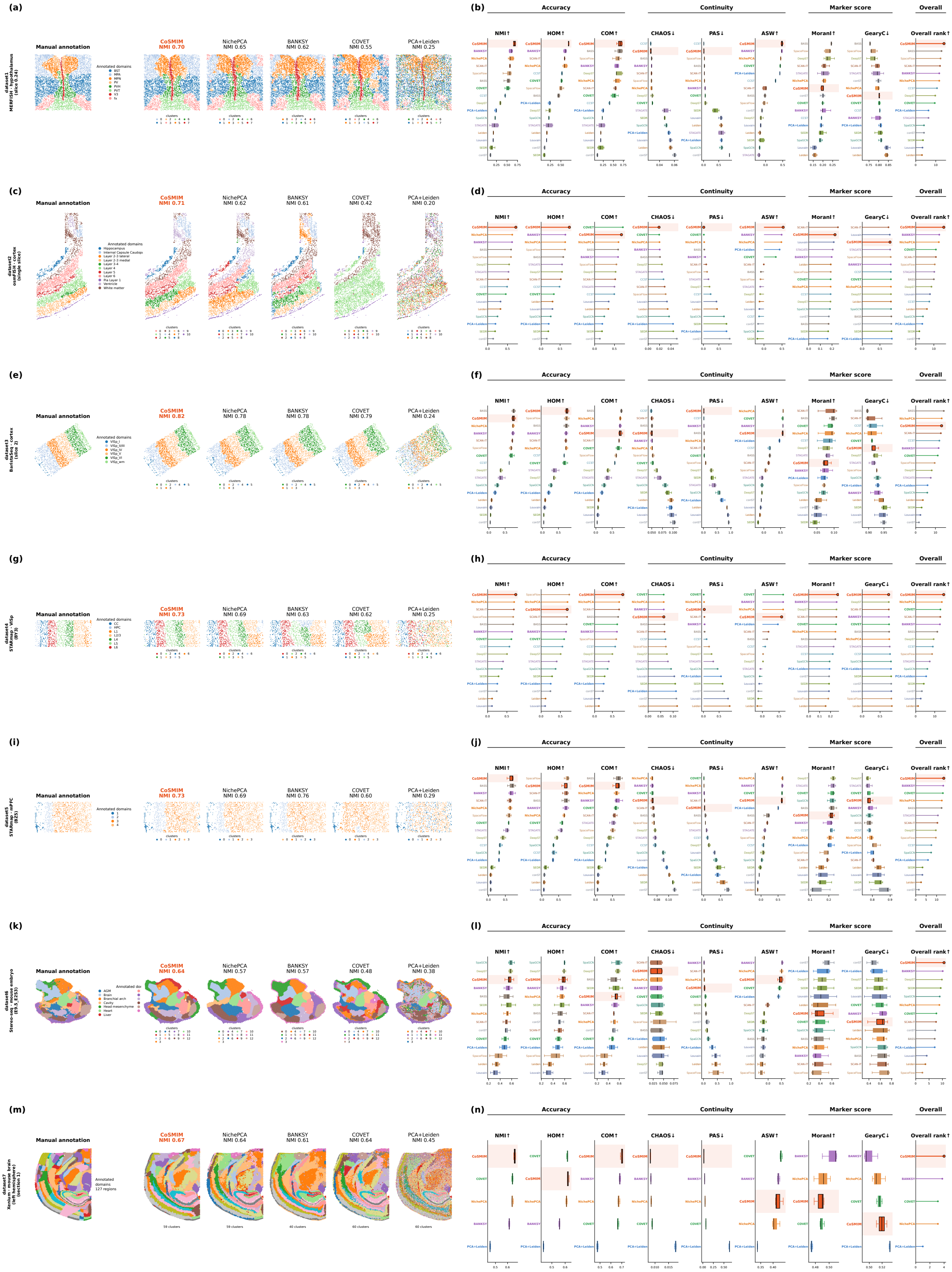


Figure S2: **Results on each of the seven datasets** (layout as in Yuan et al. 2024, Fig. 2). One row per dataset. **(a, c, e, g, i, k, m)** The annotation and the domains of CoSMIM and the four competitors on one section (named on the left), with NMI. Clusters are matched to the annotation colours. On some sections the map of a competitor comes from an earlier run at the same $k$, so the NMI printed on it can differ from Supplementary Data S1. **(b, d, f, h, j, l, n)** The eight metrics in the three groups of Section 2.4 and the overall rank (a higher rank is better), for all sections of the dataset. For dataset1 to dataset6 the published SDMBench methods are included (14 methods in total on dataset6, where SDMBench reports fewer methods). Because dataset7 is not in SDMBench, its row compares only the five methods that we ran, on the left hemisphere, with a target of 60 clusters (Section 2.5).

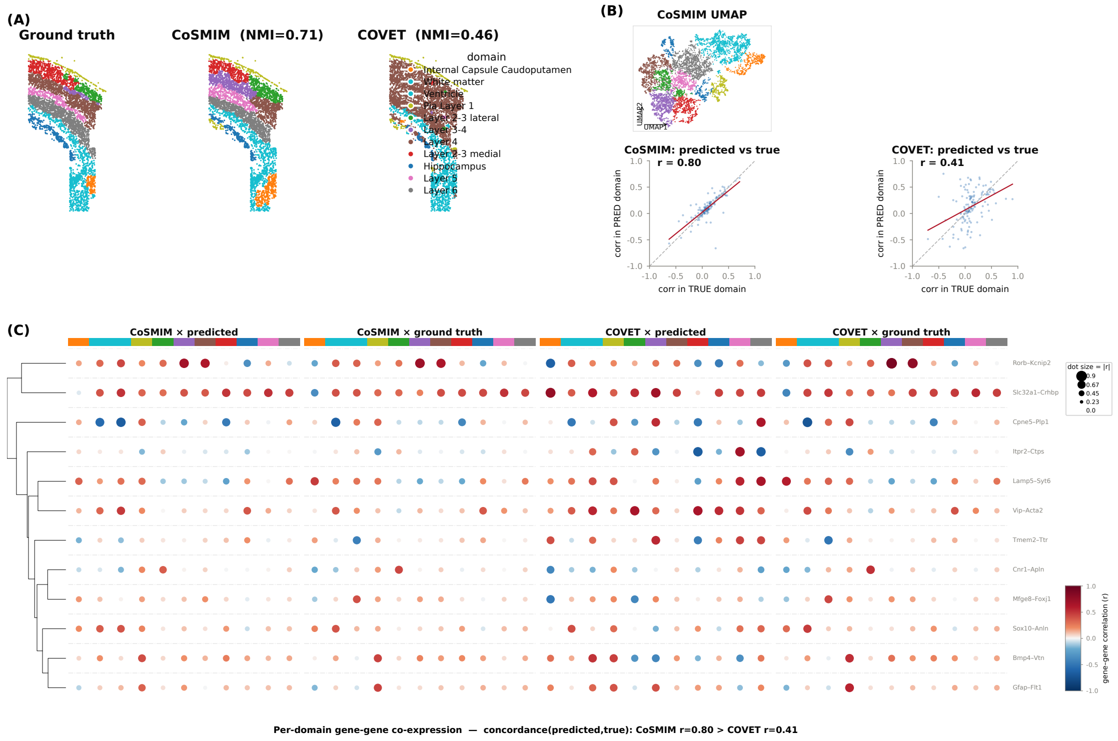


Figure S3: **Domains and per-domain correlations of CoSMIM and COVET on dataset2 (osmFISH). (A)** The annotation, CoSMIM (NMI = 0.71) and COVET (NMI = 0.46). **(B)** Top, UMAP (McInnes et al., 2018) of the CoSMIM feature. Bottom, the correlation of each of the twelve gene pairs of (C) in each predicted domain against that in the matched annotated domain: $r = 0.80$ for CoSMIM and $r = 0.41$ for COVET. Over all 496 pairs of the 32 genes, the values are 0.78 and 0.26. **(C)** The twelve gene pairs in each domain, read from the predicted and the annotated partition of each method. Dot colour shows the correlation and dot size shows its absolute value.

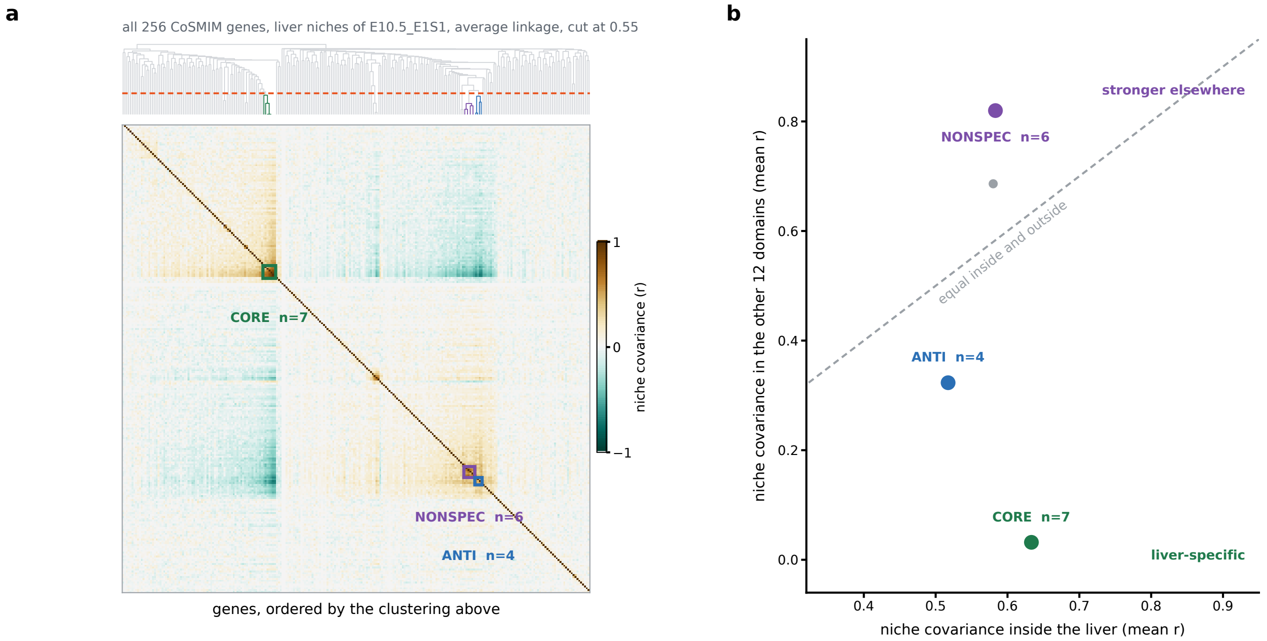


Figure S4: **Gene blocks of Fig. 4f.** **(a)** Average-linkage hierarchical clustering (distance $1 - r$) of the liver niche correlation matrix of section E10.5_E1S1 (195 liver cells, 256 genes), cut at distance 0.55 (dashed line). The matrix is ordered by the clustering. The boxes mark three blocks: CORE (7 genes), NONSPEC (6) and ANTI (4). **(b)** Every block with at least three genes: its mean correlation inside the liver (horizontal) against its mean correlation in the other twelve annotated domains of the section (vertical). Each of these domains is subsampled to at most 469 cells. The dashed line marks equal values. NONSPEC lies above this line, and CORE and ANTI lie below it. The clustering uses the liver matrix only. CORE is the liver-specific block of Fig. 4f.

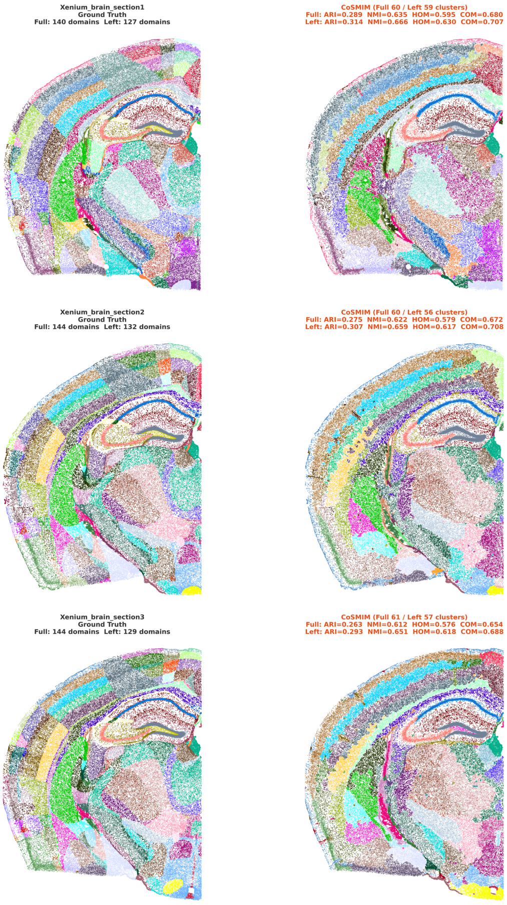


Figure S5: **Domains on dataset7 (Xenium mouse brain).** Left column, the annotation. Right column, CoSMIM. One row per section. Each panel gives the number of annotated regions or clusters, and the CoSMIM panels give ARI, NMI, HOM and COM on the whole section and on the left hemisphere. The target number of clusters is 60 (§2.5).

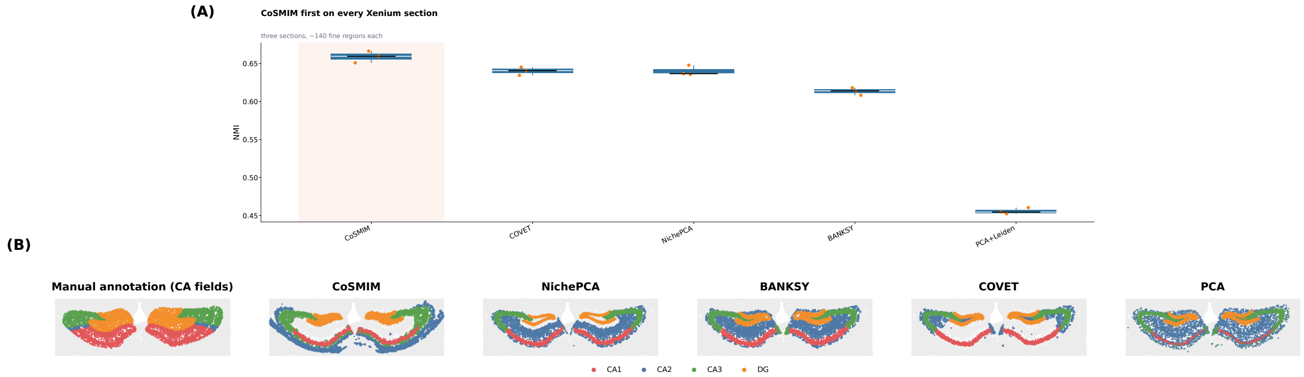


Figure S6: **Accuracy on dataset7 and the hippocampus.** **(A)** NMI of the five methods on the left hemisphere of the three sections (points). CoSMIM has the highest NMI on each section, by 0.012 to 0.026 over the best competitor. On sections 1, 2 and 3, CoSMIM has an NMI of 0.666, 0.659 and 0.651. The best competitor is COVET on section 1 (0.641) and NichePCA on sections 2 and 3 (0.648 and 0.636). The medians over the three sections are 0.659 for CoSMIM, 0.641 for COVET, 0.637 for NichePCA, 0.614 for BANKSY and 0.455 for PCA+Leiden. **(B)** The hippocampus of one section: the annotation (CA1, CA2, CA3 and DG) and the domains of the five methods.

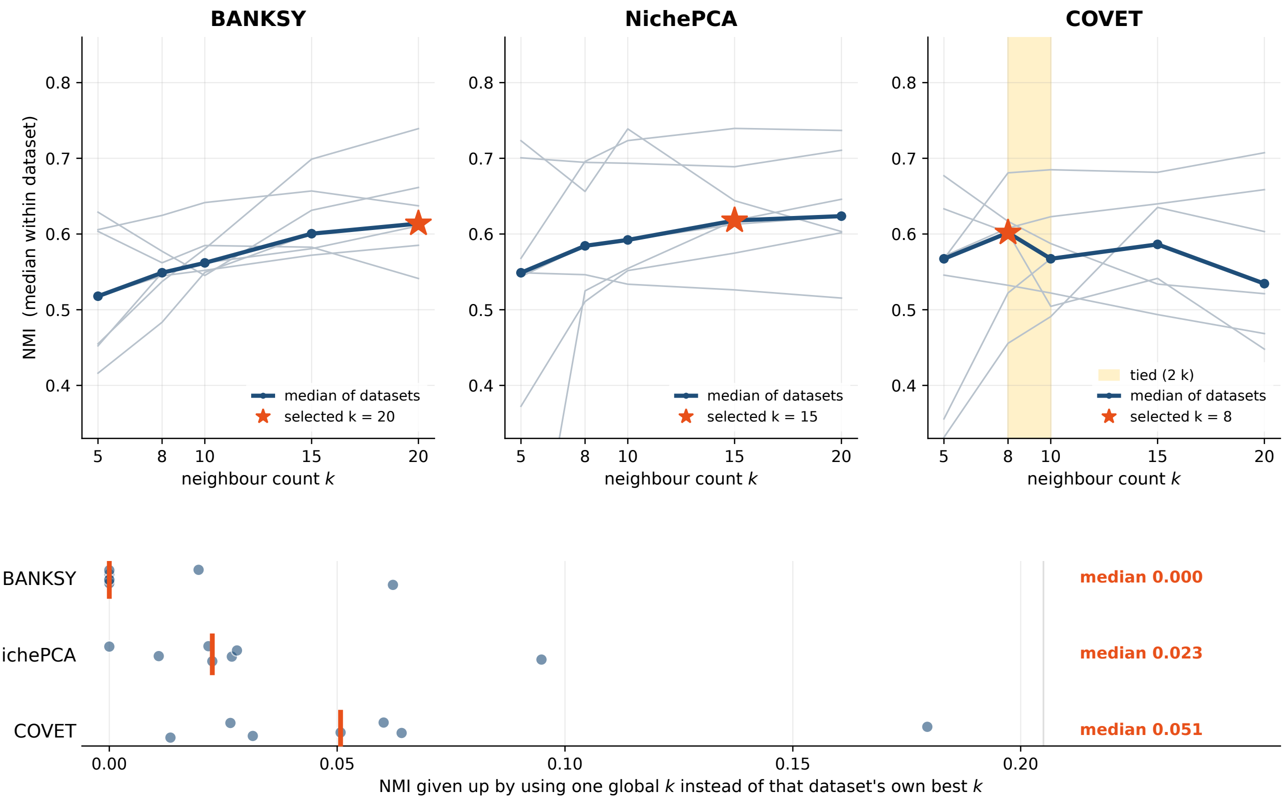


Figure S7: **Selection of the competitor neighbour count $k$.** NichePCA, BANKSY and COVET each have a neighbour count $k$. Each competitor was run at $k \in \{5, 8, 10, 15, 20\}$ on dataset1 to dataset6 and ranked against the eleven published SDMBench methods, one $k$ at a time. For each method we selected the $k$ with the best mean rank over NMI, HOM and COM. A tie is broken by the higher mean of NMI, HOM and COM. This gives NichePCA $k = 15$, BANKSY $k = 20$ and COVET $k = 8$, and this $k$ is used on all seven datasets. dataset4 is scored against the SDMBench annotation. **Top**: NMI of BANKSY, NichePCA and COVET at each $k$. Grey lines are the medians over the sections of each of the seven datasets. The blue line is their median. The star marks the selected $k$. The shaded band marks the $k$ values with the same accuracy rank as the selected $k$. dataset7 is shown on whole sections and is not used for the selection. **Bottom**: for each dataset (points), the NMI lost by using the selected $k$ instead of the best $k$ for that dataset. The bar marks the median. In the benchmark, COVET uses at most 64 highly variable genes, the default of its implementation. NichePCA and BANKSY use every gene that passes quality control. NichePCA uses 8 principal components and a clustering graph with $k$ neighbours. BANKSY uses a mixing weight of 0.8 and 15 principal components. COVET and PCA+Leiden use up to 30 principal components and 50 graph neighbours. For every method, the Leiden resolution is searched for the number of clusters closest to the number of annotated domains, or to 60 on dataset7 (Section 2.5). The ranks and the values at each $k$ are in Supplementary Data S1.

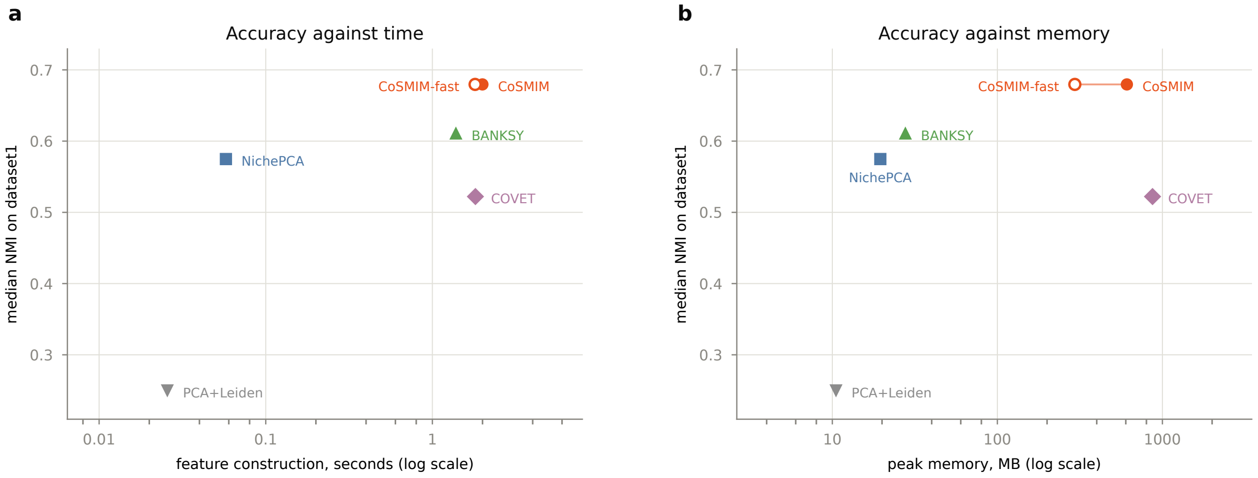


Figure S8: **Cost and accuracy of the five methods on dataset1.** CoSMIM has no training stage, so its cost is the construction of the feature: $O(N\,m\,k_m\,g^2)$ operations for the covariances of $N$ cells at $m$ scales and $O(Ng^3)$ for the matrix square roots. **(a)** Median wall time of feature construction against median NMI over the five dataset1 sections, on one node, with each competitor at its benchmark $k$: 2.00 s for CoSMIM, 1.81 s for COVET, 1.38 s for BANKSY, 0.06 s for NichePCA and 0.03 s for PCA+Leiden. The shared downstream steps (PCA, $k$NN graph and Leiden) are not included. **(b)** Peak memory of the same runs against the same NMI: 609 MB for CoSMIM, 871 MB for COVET, 28 MB for BANKSY and 20 MB for NichePCA. A covariance stores $g(g+1)/2$ numbers per cell and a mean stores $g$. CoSMIM-fast is the single-precision version of the same feature. It differs from the double-precision feature by a relative Frobenius distance of about $10^{-7}$ and needs 295 MB. All results in this paper use double precision.
